# MAAPER: model-based analysis of alternative polyadenylation using 3’ end-linked reads

**DOI:** 10.1101/2021.03.21.436343

**Authors:** Wei Vivian Li, Dinghai Zheng, Ruijia Wang, Bin Tian

## Abstract

Most eukaryotic genes harbor multiple cleavage and polyadenylation sites (PASs), leading to expression of alternative polyadenylation (APA) isoforms. APA regulation has been implicated in a diverse array of physiological and pathological conditions. While RNA sequencing tools that generate reads containing the PAS, named *onSite* reads, have been instrumental in identifying PASs, they have not been widely used. By contrast, a growing number of methods generate reads that are close to the PAS, named *nearSite* reads, including the 3’ end counting strategy commonly used in single cell analysis. How these nearSite reads can be used for APA analysis, however, is poorly studied. Here, we present a computational method, named model-based analysis of alternative polyadenylation using 3’ end-linked reads (MAAPER), to examine APA using nearSite reads. MAAPER uses a probabilistic model to predict PASs for nearSite reads with high accuracy and sensitivity, and examines different types of APA events, including those in 3’UTRs and introns, with robust statistics. We show MAAPER’s accuracy with data from both bulk and single cell RNA samples and its applicability in unpaired or paired experimental designs. Our result also highlights the importance of using well annotated PASs for nearSite read analysis.

## Introduction

Almost all eukaryotic mRNA genes employ cleavage and polyadenylation (CPA) for 3’ end processing (Proudfoot, 2016). Well over half of the genes harbor multiple PASs, resulting in expression of alternative polyadenylation (APA) isoforms (Gruber and Zavolan, 2019; Tian and Manley, 2017). While most of the APA events occur in the 3’-most exon of genes, leading to isoforms with variable 3’ untranslated regions (3’UTRs), a sizable fraction of APA sites, e.g., ∼20% in the human genome, are located in introns, the usage of which additionally leads to alternation of coding sequence (CDS). APA is increasingly being appreciated as a major mechanism for gene regulation (Gruber and Zavolan, 2019; Tian and Manley, 2017), diversifying the transcriptome in different cell types and under various pathological and physiological conditions. To accurately profile APA isoforms is of great importance in understanding the mechanisms and consequences of APA.

The advent of RNA-seq technologies has enabled comprehensive transcriptome analysis. RNA-seq data have also been used to examine APA isoform profiles, either by taking advantage of drop of RNA-seq read coverage at the PAS (Xia et al., 2014) or by using annotated PASs (Grassi et al., 2016; Ha et al., 2018; Wang and Tian, 2020). However, because RNA-seq data are not designed to identify PASs, these approaches lack high accuracy and sensitivity for PAS identification and APA profiling.

RNA sequencing methods that generate reads biased to the 3’ end of transcripts, collectively called 3’ end sequencing here, are increasingly being used to study gene expression in bulk RNA samples or in single cells (Zhang et al., 2019). Some methods generate reads that end at a PAS, such as PAS-seq (Shepard et al., 2011), SAPAS (Fu et al., 2011), MAPS (Fox-Walsh et al., 2011), and PolyA-seq (Derti et al., 2012). For simplicity, we name those reads *onSite* reads. As such, the PAS can be directly identified after mapping the reads to the reference genome (Figure 1A). While straightforward for PAS identification, these methods typically require a custom primer containing an oligo(T) region for sequencing (Moll et al., 2014), limiting their applications. By contrast, some methods generate reads that are close to but do not necessarily contain the PAS, such as QuantSeq FWD (Moll et al., 2014) and PAT-seq (Harrison et al., 2015) (Figure 1A). For simplicity, those reads are named *nearSite* reads, Conceivably, nearSite reads are “3’ end-linked”, because they embody information about a PAS. Notably, several 3’ end counting protocols used in single-cell RNA sequencing (scRNA-seq), such as 10x Genomics (Zheng et al., 2017) and Drop-Seq (Macosko et al., 2015), also generate nearSite reads (Patrick et al., 2020; Shulman and Elkon, 2019). As such, nearSite read data have grown rapidly in recent years. However, mining nearSite reads for APA has hitherto been limited due to lack of suitable methods. The major challenge of using nearSite reads for APA analysis is accurate assignment of reads to a PAS. Here we present a computational method named MAAPER (<u>m</u>odel-based <u>a</u>nalysis of <u>a</u>lternative <u>p</u>olyadenylation using 3’ <u>e</u>nd-linked <u>r</u>eads), which accurately assigns nearSite reads to PASs through statistical modeling, and generates multiple statistics for APA analysis.

**Figure 1.**
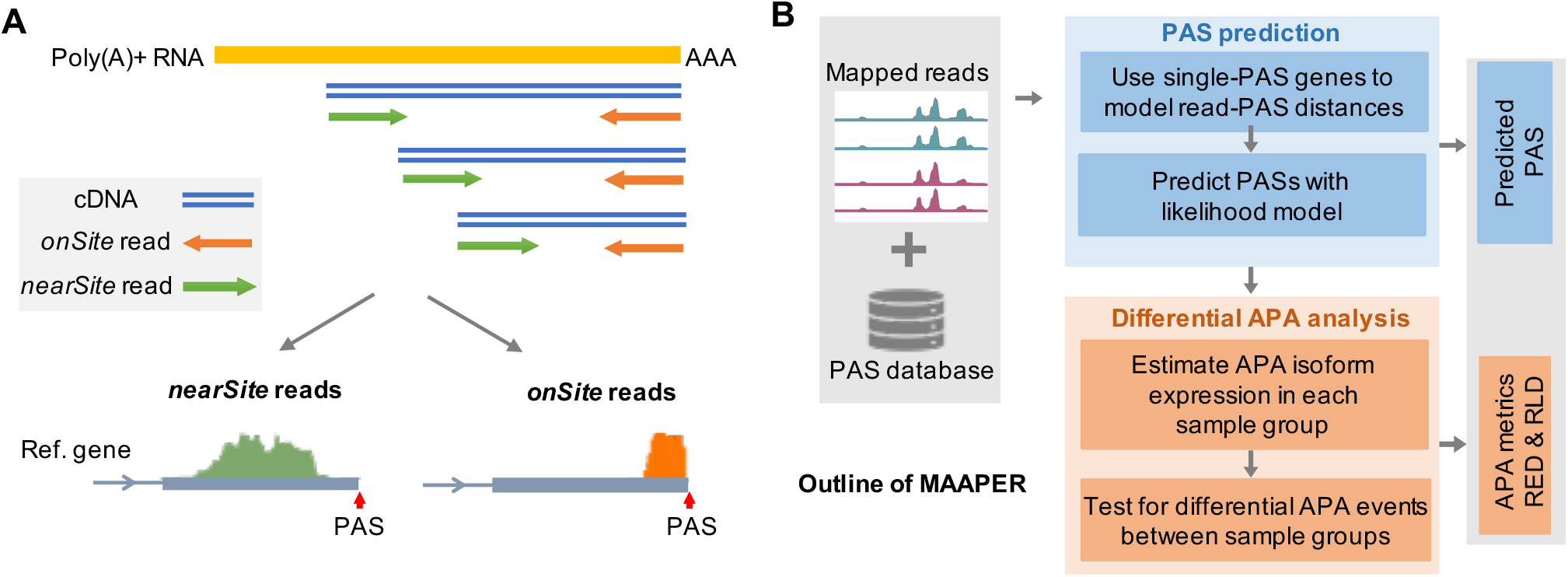
Overview of the MAAPER method. (**A**) A schematic illustrating nearSite reads (e.g., QuantSeq FWD, 10x Genomics) and onSite reads (e.g., QuantSeq REV). (**B**) Outline of the MAAPER method. MAAPER takes aligned nearSite reads and a PAS database (polyA_DB) and outputs predicted PAS positions and APA analysis results.

## Results and Discussion

### Design of MAAPER

While nearSite reads do not directly map to PASs, they have implicit information about the PASs that give rise to the reads. We reasoned that the distance between the 5’ end of a nearSite read and PAS reflects the cDNA fragment size of a sequencing library and thus could be statistically modelled. We further reasoned that genes that contain only a single PAS would provide a clear picture for studying the read-PAS relationship, and the information derived can be used to study genes containing multiple PASs. To these ends, we developed a strategy to first predict PASs (stage 1) and then use them for APA analysis (stage 2). We named our method MAAPER (model-based analysis of alternative polyadenylation using 3’ end-linked reads, Figure 1B).

At the PAS prediction stage, MAAPER first uses genes that contain only a single PAS to learn the distribution of read-PAS distance (Figure 1B). PolyA_DB database, a comprehensive PAS database our lab has constructed (Wang et al., 2018), is used to identify those genes (Figure S1). It is worth noting that the current version (v3) of PolyA_DB contains PASs mapped by 3’ end extraction and sequencing (3’READS), a highly accurate method for PAS identification (Hoque et al., 2013). The statistical learning is based on a probabilistic model that employs the Gaussian kernel function for non-parametric estimation of read-PAS distances. This approach captures the distributional characteristic of read-PAS distances without assuming data uniformness. Next, MAAPER uses the learned read-PAS relationship and a likelihood model to predict PASs for each gene given its mapped nearSite reads. Each nearSite read has a set of probability scores for all the annotated PASs of a given gene (Figure S2A), and these scores indicate the likelihood of the PASs. Again, PAS information in the PolyA_DB data is used to guide the prediction for high accuracy.

At the APA analysis stage, MAAPER uses the predicted PASs for quantitative analysis of APA difference between samples (Figure 1B). For each gene, the expression of a PAS is based on the proportion of RNA isoforms containing the site in a sample. Three statistical tests can be carried out by MAPPER. Given unpaired experimental designs, MAAPER uses a likelihood ratio test (LRT) to determine if a gene’s overall PAS profile differs between conditions and a Fisher’s exact test to determine if two selected PASs’ relative abundance differs between conditions. Given paired experimental designs, MAAPER uses an *F* test to account for the correlation structures and detects consistent APA changes between conditions (see Methods).

### Accurate and sensitive PAS prediction by MAAPER

To aid in MAAPER development, we used the QuantSeq FWD method to generate nearSite data from mouse NIH3T3 cells under four conditions, namely, nontreated cells (NT), cells treated with arsenite stress (AS), and cells that have recovered from AS treatment for 4 or 8 hours (RC4 or RC8) (Figure 2A, see Methods for detail). We chose these four conditions because we previously found substantial gene expression and APA changes across these conditions (Zheng et al., 2018). We generated ∼28M mapped reads per sample (Table S1). For comparison and validation, we subjected the same RNA samples to sequencing by QuantSeq REV, a method that generates onSite reads. Importantly, QuantSeq FWD and REV methods have the same protocols except for the sequencing step, allowing us to compare nearSite reads and onSite reads more accurately without technical noise coming from sample acquisition or processing.

**Figure 2.**
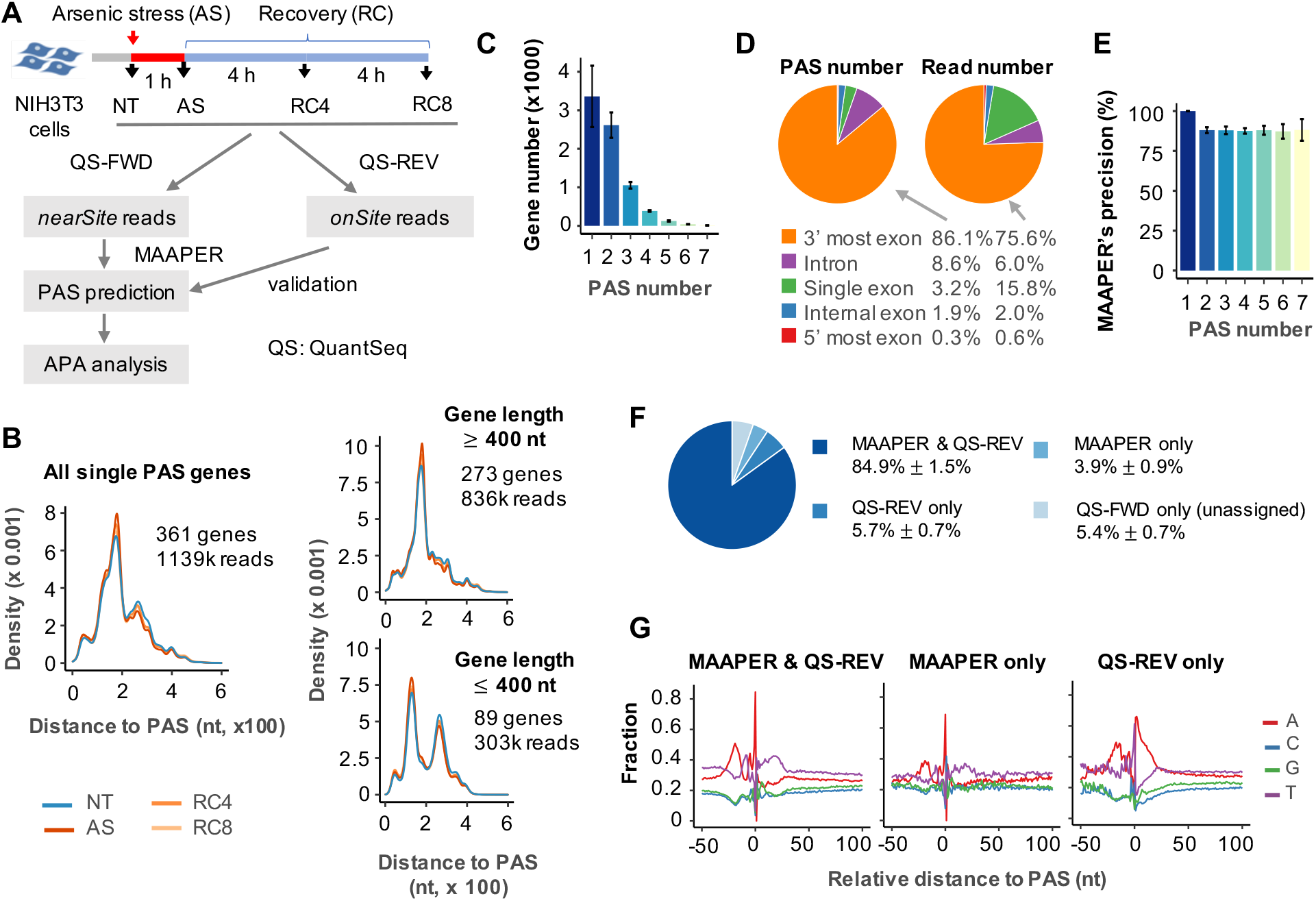
Accurate and sensitive PAS prediction by MAAPER. (**A**) Design of experiment using QuantSeq FWD (QS-FWD) and QuantSeq REV (QS-REV) to study APA in NIH3T3 cells treated by arsenic stress (AS). (**B**) Distribution of read-PAS distances based on genes with a single PAS. (**C**) Number of genes with a varying number of predicted PASs by MAAPER from QuantSeq FWD data. Bars are the average gene number across all samples, and Error bars are SD. (**D**) Proportion of predicted PASs and FWD reads based on their genomic positions. Proportions were averaged across all samples. (**E**) Precision rates of MAAPER’s PAS prediction for genes with a varying number of PASs. Bars are the average precision across all samples. Error bars are SD. (**F**) Proportions of four categories of QuantSeq FWD reads. Mean and SD of read proportions are based on all samples. (**G**) Nucleotide frequencies around three categories of predicted PASs.

Using the nearSite reads from QuantSeq FWD, we detected 7,870 genes on average from the NT and AS-treated samples. Of these, 361 were annotated as single-PAS genes in the PolyA_DB database (Wang et al., 2018), which we then focused on for modeling of the read-PAS distance. Interestingly, we found that read-PAS distance had two modes, one centered around 190 nt from the PAS and another 250 nt from the PAS. Further analysis indicated that the 250 nt peak came from short genes (<400 nt) (Figure 2B). Therefore, we only used the first peak to represent read-PAS distribution in our analysis.

On average, 44.3% of detected genes used a single PAS in the four samples and 34.4% two PASs, 13.9% three PASs, and 7.4% >3 PASs (Figure 2C). These numbers are quite consistent across the four samples, with a standard deviation (SD) of 3.9%, 1.1%, 1.3%, and 0.4%, respectively. We analyzed the locations of the predicted PASs and found that 89.3% PASs were in the 3’ most exons (including single-exon genes) and 8.6% in introns (Figure 2D, left). These corresponded to 91.4% and 6.0%, respectively, in terms of read number (Figure 2D, right).

To evaluate our result, we used the onSite reads from QuantSeq REV to identify PASs (based on the 3’-most position of each read). Because data were generated from the same samples, the onSite reads practically served as an *ad hoc* PAS database in the same biological context, in lieu of PolyA_DB. With the 11M mapped onSite reads from QuantSeq REV (Table S1), we identified ∼16K PASs in the four samples. Based on these PASs, we calculated precision and recall rates of MAAPER, with precision being defined as the percentage of PASs identified by nearSite reads that matched those by onSite reads, and recall as the percentage of PASs identified by onSite reads that matched those by nearSite reads. MAAPER achieved an overall precision of 90.7% (SD = 2.0%) across the four samples, and 87.2%-100% for genes with different numbers of PAS (Figure 2E); the overall recall for all genes was 75.5% (SD = 2.5%), and 81.9% (SD = 1.7%) for genes with ≤3 PASs (Figure S3A-C).

Interestingly, we found that while the PASs that were identified both by onSite reads and predicted by MAAPER (“MAAPER & QS-REV”, 84.9% of all PASs) or PASs that were identified by MAAPER only (“MAAPER only”, 3.9% of all PASs) showed similar nucleotide profiles around the PAS (Figures 2F-G and S4), with upstream A-rich and downstream U-rich peaks that are characteristic of PASs (Tian and Graber, 2012), the PASs that were identified by onSite reads but not by MAAPER (“QS-REV only”, 5.7% of all PASs) showed A-rich peaks both in upstream and downstream regions (Figures 2G and S4). This result indicates that “QS-REV only” reads were likely to have been generated by priming of oligo(dT) at internal A-rich sequences instead of the poly(A) tail, a problem known as *internal priming*. The internal priming issue is commonly associated with methods using oligo(dT) for cDNA construction (Nam et al., 2002), but is well addressed in methods that do not depend on oligo(dT), such as 3’READS (Hoque et al., 2013). Because the PASs annotated in the PolyA_DB database v3 were based on the 3’READS method only, the PASs predicted by MAAPER are not affected by the internal priming issue (Figure 2G). As expected, MAAPER achieved a higher overall recall of 86.6% (SD = 2.0%) after the “QS-REV only” reads were excluded (Figure S3D-F).

### Differential APA analysis by MAAPER

For differential analysis of APA, MAAPER employs likelihood ratio tests to determine the statistical significance of observed differences in isoform abundance between samples. Two scores are generated, namely, <u>r</u>elative <u>l</u>ength <u>d</u>ifference (RLD) and <u>r</u>elative <u>e</u>xpression difference (RED) (Figure S2B; see Methods for detail). RLD*u* measures the relative 3’UTR size changes based on all PASs in the 3’-most exon, whereas RLD*i* measures the relative pre-mRNA size when there are intronic PASs. Positive RLD*u* or RLD*i* values indicate increased expression of isoforms using distal PASs, resulting in 3’UTR or pre-mRNA lengthening, respectively. The RED score is based on the top two most differentially expressed isoforms (distal PAS isoform/proximal PAS isoform, log2 ratio), regardless of their locations. As such, a positive RED score indicates increased expression of the distal PAS isoform and a negative RED score increased expression of the proximal isoform.

Applying MAAPER to the QuantSeq FWD data, we examined the statistical significance of APA between AS, RC4, or RC8 and NT samples. The overall distributions of RED and RLD scores indicated stronger APA regulation in RC4 and RC8 samples compared with the AS sample (Figure S5A). Using FDR-adjusted *p*-value ≤ 0.01 as the cutoff, we identified significant APA events in the AS-treated samples (Figure 3A). Consistent with our previous finding, AS, RC4, and RC8 samples all displayed preferential expression of proximal PAS isoforms (median RED = -0.31, -0.62, and -0.21, respectively, Figure 3B). Based on RLD*u*, 3’UTRs shortened after stress, with the RC4 sample showing the greatest extent (median RLD*u* = -0.07, -0.14, and -0.05 for AS, RC4, and RC8, respectively, Figure 3B). Several example genes, including *Nmt1, Calm1, Timp2*, and *Purb*, are shown in Figures 3C and S6A. Their MAAPER results were completely consistent with our previous results using 3’READS and validated by RT-qPCR (Zheng et al., 2018).

**Figure 3.**
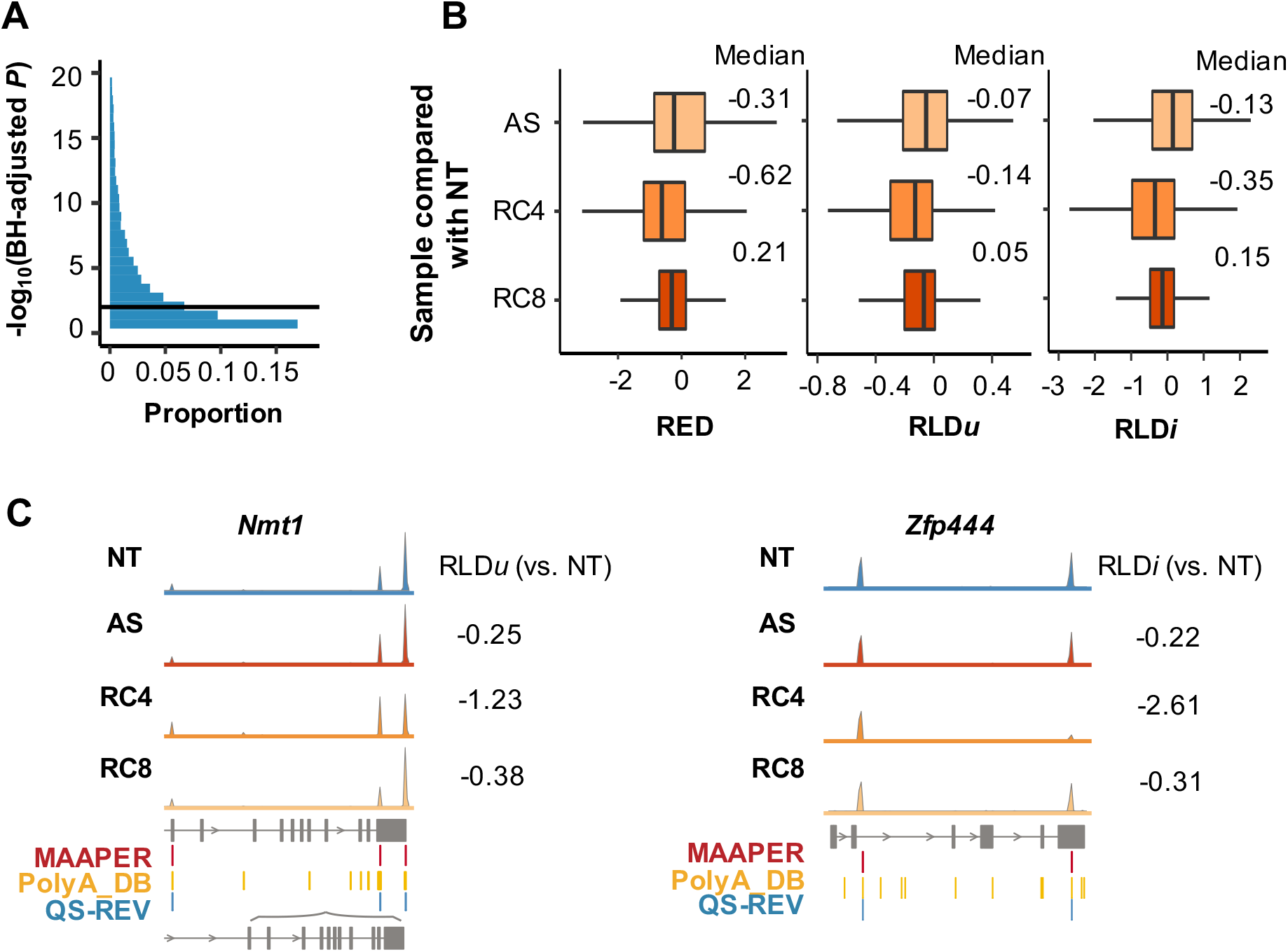
Differential APA analysis by MAAPER on QuantSeq FWD data. (**A**) Distribution of -log10(FDR-adjusted *p* value) in differential APA analysis. Data were based on average across all samples. The threshold of adjusted *p* value = 0.01 was used to identify significant changes between AS or RC4 or RC8 and NT. (**B**) Boxplots of RED, RLD*u*, and RLD*i* scores calculated for AS, RC4, and RC8 samples. Median values are labelled on the right. Only genes that showed significant APA changes in at least one sample compared with NT were included. (**C**) Example genes *Nmt1* and *Zfp444*.

Notably, based on RLD*i*, MAAPER revealed global pre-mRNA shortening through intronic polyadenylation in AS and RC samples (median RLD*i* = -0.13, -0.35, and 0.15 in AS, RC4, and RC8 samples, respectively, Figure 3B). This was not reported before, and indicates that stress shortens both 3’UTRs and pre-mRNAs. Several example genes, including *Zfp444, Taf6, Nup155, and Elf1*, are shown in Figures 3C and S6B. Gene ontology (GO) enrichment analysis showed that, genes with significant 3’UTR shortening (negative RLD*u* scores) in the RC4 sample tended to encode proteins involved in amide metabolic processes, RNA or protein catabolic processes, and RNA stability (Figures S7). In contrast, genes with significant pre-mRNA shortening (negative RLD*i* scores) tended to encode proteins involved in telomerase activity and telomere organization, RNA splicing, and mRNA processing (Figures S8).

### MAAPER enables APA analysis with single cell data

A growing number of scRNA-seq experiments generate nearSite reads (Zhang et al., 2019; Zheng et al., 2017). We next wanted to use MAAPER to analyze APA events in single cell populations. To this end, we downloaded 10X Genomics data of placental samples (Vento-Tormo et al., 2018), which included scRNA-seq data from five donors (D8-D12). We focused on the three trophoblast cell populations we recently found to have substantial APA differences (Cheng et al., 2020), namely, villous cytotrophoblasts (VCTs), extravillous trophoblasts (EVTs) and syncytiotrophoblasts (SCTs) (Figures 4A and S9). Because VCTs are the precursor cells of the other two subtypes (Chang et al., 2018), we compared APA profiles of EVTs of SCTs with those of VCTs.

**Figure 4.**
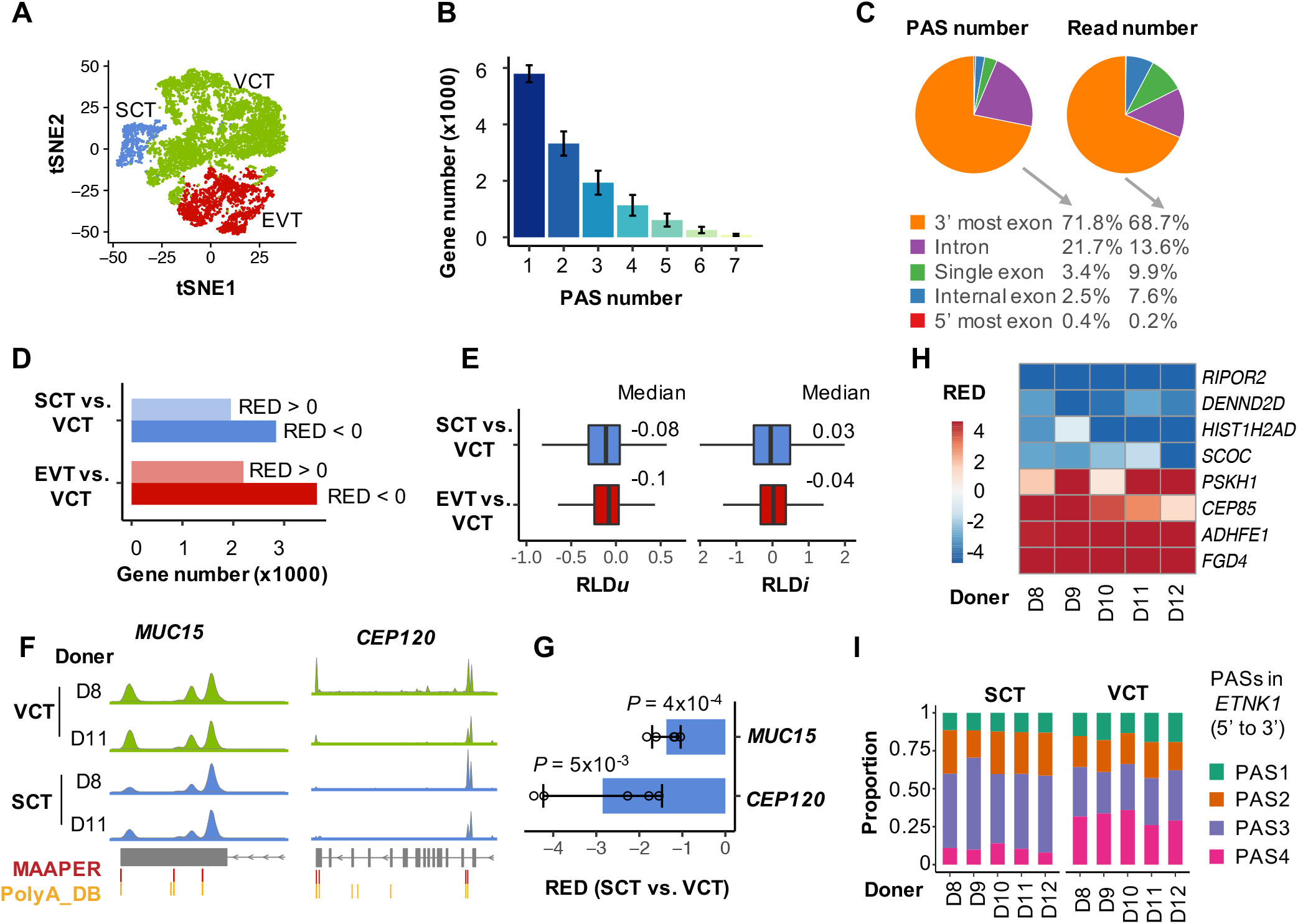
Differential APA analysis by MAAPER on 10x Genomics data. (**A**) tSNE plot of the three trophoblast cell types based on gene expression. (**B**) Number of genes with a varying number of predicted PASs based on the scRNA-seq data. Bars are the average gene number across three cell types; Error bars are SD. (**C**) Proportion of predicted PASs and FWD reads based on their genomic positions. Proportions were averaged across all samples. (**D**) Number of genes with significant APA changes in EVTs or SCTs in comparison with VCTs. Genes are categorized according to the signs of their RED scores. (**E**) Boxplots of RLD*u* and RLD*i* scores calculated for EVTs and SCTs, based on genes that showed significant APA changes in each cell type. Median values are labelled on the right. (**F**) Example genes *MUC15* and *CEP120*. Aggregated reads from two donors (D8 and D11) are displayed as examples. (**G**) RED scores of *MUC15* and *CEP120* for all donors. The *p* value of one-sided t test is 0.0004 for *MUC15* and 0.005 for *CEP120*. (**H**) RED scores of eight example genes with significant APA changes identified by the paired test between SCTs and VCTs. (**I**) PAS proportions of the *ETNK1* gene in SCTs and VCTs (adjusted *p* = 0.05 in paired test).

MAAPER on average identified 13,156 expressed genes in the three trophoblast cell populations, among which 44.0% had a single PAS, 25.2% two PASs, 14.7% three PASs, and 16.1% >3 PASs (Figure 4B). We found that 75.2% PASs were in the 3’-most exons (including single-exon genes) and 21.7% were in introns, accounting for 78.6% and 13.6% of all reads, respectively (Figure 4C).

Using FDR-adjusted *p*-value ≤ 0.01 as the cutoff, MAAPER detected significant PAS changes in 5,843 genes between EVTs and VCTs and 4,796 genes between SCTs and VCTs (Figure 4D). The median RED was -0.35 for EVTs vs. VCTs and -0.48 for SCTs vs. VCTs, indicating increased abundance of proximal PAS isoforms in both EVTs and SCTs (Figure S10A). Interestingly, the genes with significant APA changes in EVTs or SCTs did not overlap substantially (Figure S10B), indicating distinct APA regulation.

The median RLD*u* score were -0.08 and -0.10 for EVTs vs. VCTs, and SCTs vs. VCTs respectively (genes with significant APA only), indicating global shortening of 3’UTRs (Figure 4E). For example, MAAPER predicted three PASs in the 3’UTR of *MUC15*, which encodes for membrane-bound mucins and acts as a negative regulator of trophoblast invasion (Shyu et al., 2007). We found that *MUC15* was not only differentially expressed between the three cell types (Figure S11A), but also displayed 3’UTR shortening in the differentiation of SCTs (FDR-adjusted *p* value ≈ 0; RLD*u* = -0.71; Figures 4F and S11B). RED and RLD*u* scores of all donors are shown in Figures 4G and S11C, both of which were significantly below zero.

The median RLD*i* scores were 0.04 in EVTs vs. VCTs and -0.03 in SCTs vs. VCTs, respectively (genes with significant APA only), indicating distinct intronic APA regulations in differentiation of SCTs compared to that of VCTs (Figure 4E). For example, MAAPER identified four PASs on the *CEP120* gene, which encodes for a protein that functions in the microtubule-dependent coupling of the nucleus and the centrosome (Figure 4F and S11D-E). Two of the PASs were located in the 3’ most exon and another two were in an intron. MAAPER detected strong intronic polyadenylation activation in SCTs compared to VCTs (FDR-adjusted *p* value ≈ 5.3 × 10^−48^; RLD*i* = -2.27; Figures 4F). The estimated proportion of transcripts using this intronic PAS increased from 53.9% in VCTs to 87.5% in SCTs. In addition, RED and RLD*i* scores of all donors are shown in Figures 4G and S11F, both of which were significantly below zero.

Single cells from a given sample are naturally paired by sample identity. In the case of placental samples, distinct cell populations could be compared within each individual and their differences across individuals could be analyzed to derive inter-sample variation. To this end, we designed a paired test to detect differential APA events that were consistent across individuals (see Methods for details). Using this paired test, we identified 138 genes with significant APA changes between SCTs and VCTs and 182 genes between EVTs and VCTs (FDR-adjusted *p* value < 0.1). As expected, these genes consistently showed transcript lengthening or shortening across the five donors (Figures 4H and S12). For example, MAAPER identified four PASs in the *ETNK1* gene, which repeatedly displayed 3’UTR shortening in SCTs (Figure 4I). In contrast, significant genes identified by the unpaired test did not always present consistent changes among donors (Figure S14). When focusing on genes with highly variable APA profiles in SCTs, we found that a subset of three donors (D8, D10, and D11) demonstrated higher similarity (Figure S15).

In summary, MAAPER is the first method that predicts PASs and conducts APA analysis using nearSite reads. Unlike most existing RNA-seq-based tools that model only two PASs, MAAPER predicts a comprehensive set of APA isoforms for a gene, including those using PASs located in 3’ UTRs and introns. By incorporating the large collection of well annotated PASs in PolyA_DB database (Wang et al., 2018), MAAPER identifies PASs with high accuracy and sensitivity. With a rigorous statistical framework based on a likelihood model, MAAPER provides statistically interpretable results as well as multiple metrics indicating the extent and consequences of APA regulation. Paired APA analysis, first in its class, identifies common and distinct APA events in cell populations from multiple individuals.

## Materials and Methods

### Cell culture and RNA sequencing

NIH3T3 cells were cultured in high glucose Dulbecco’s modified Eagle’s medium (DMEM) supplemented with 10% calf serum, 100 IU/ml penicillin, and 100 μg/ml streptomycin at 37 °C. For stress induction, cells were cultured in medium containing 250 μM of sodium arsenite (NaASO2) for 1 h. For recovery, stressed cells were washed twice with phosphate-buffered saline (PBS) to remove NaASO2, followed by culturing in regular medium for 4 h or 8 h. Total RNA was extracted using TRIzol. QuantSeq 3’ mRNA-Seq Library Prep Kits (FWD or REV) were used to prepare libraries. Sequencing was carried out on an Illumina HiSeq.

### Summary of the MAAPER method

MAAPER is the first method for predicting PAS locations and comparing APA usage from nearSite reads, and it has four unique features which contribute to its high accuracy and broad applicability. First, the statistical model of MAAPER is flexible. Unlike most existing PAS prediction tools (develop for RNA-seq) that use a simplified model of only two PASs (proximal and distal), MAAPER predicts a comprehensive set of expressed PASs for each gene, including those located in 3’ UTRs, internal exons, and introns. Second, MAAPER has high accuracy in PAS prediction. On one hand, it achieves a high recall rate by incorporating the large collection of PASs annotated in PolyA_DB database, which contains more than 85,000 and 120,000 PASs for human and mouse genes in this current version (v3). On the other hand, it achieves a high precision by a forward-selection procedure, which screens for a conservative set of PASs that adequately explain the observed reads. Third, MAAPER is statistically principled. It detects differential PAS usage between conditions using rigorous and flexible statistical tests. Most of the aforementioned methods for RNA-seq or onSite reads use the Fisher’s exact test (Fisher, 1922) to test for differential PAS usage, which is limited to comparisons between two samples and two PASs (Grassi et al., 2016; Ha et al., 2018; Wang and Tian, 2020; Xia et al., 2014). In contrast, for unpaired samples, MAAPER formulates an LRT based on its likelihood model to accurately detect if a gene’s PAS proportions change between conditions; for paired samples, MAAPER uses an *F*-test to account for the correlation structures and detects consistent APA changes between conditions. Fourth, MAAPER facilitates the interpretation of differential analysis of PAS usage. In addition to estimated PAS proportions, it calculates two scores to measure the differences in relative transcript length and relative PAS expression, providing a convenient approach to quantifying the strength of APA regulation signals.

### Likelihood model for PAS prediction

MAAPER uses a rigorous and flexible likelihood-based framework to identify PASs and quantify APA isoform abundance with 3’ end-linked RNA sequencing data (nearSite reads). There are two steps at the PAS prediction stage. At the first step, MAAPER learns the distance of nearSite reads to PASs using only genes with a single PAS. At the second step, MAAPER uses a likelihood model to predict PASs for all reads. MAAPER uses data from PolyA_DB database (v3), a comprehensive PAS database based on 3’READS data (Wang et al., 2018), at both steps. Specifically, for each sample, after aligning reads to the reference genome, we extract all the reads mapped to the single PAS genes (annotated in PolyA_DB) and calculate the distances (excluding intronic regions) between the 5’ positions of these reads and their corresponding PASs. Next, we non-parametrically estimate the probability density of the distances using a Gaussian kernel function. This is achieved with the density function in R. This approach allows us to capture the distributional characteristic of the distances without assuming uniformness, which is often violated in real data (Li et al., 2019). In our analysis, we learn distances for genes ≤ 400 nt and genes > 400 nt separately, as we have found that the distances for short genes displayed different distributional properties (Figure 2B).

At the second step, MAAPER constructs a likelihood model to identify PASs and quantify isoform abundance. Suppose a given gene has *J* annotated PASs in PolyA_DB, then we denote their proportions in the mRNAs transcribed from this gene as 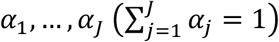 . Note that the model aims to detect all PASs in all samples. For sample replicates, we denote the replicate number as *K*_1_ in condition 1 and as *K*_2_ in condition 2. When there is no replicate, *K*_1_ = *K*_2_ = 1. In addition, we use *n*_*ck*_ to denote the number of reads mapped to a gene in the *k*-th replicate in condition *c* (*c* = 1 or 2). Given the above notations, the joint likelihood of RNA-seq reads mapped to this gene can be derived as

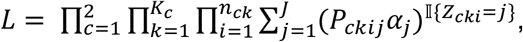

where *Z*_*cki*_ ∈ {1, …, *J*} is the unobserved PAS index and *Z*_*cki*_ = *j* if the *i*-th read in the *k*-th replicate in condition *c* comes from a transcript using the *j*-th PAS. *P*_*ckij*_ denotes the probability of observing a read at read *i*’s 5’ position in the *k*-th replicate in condition *c*, given that the read comes from a transcript using the *j*-th PAS; *P*_*ckij*_ is calculated based on the distance densities learned from the first step.

Given the above likelihood function, we developed an Expectation-Maximization (EM) algorithm (Dempster et al., 1977) to maximize the log-likelihood

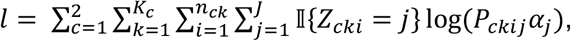

and estimate the PAS proportions. We use 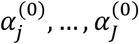 to denote initial values and use 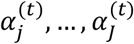 to denote estimated values in the *t*-th iteration of the EM algorithm. Then, in the (*t* + 1)-th iteration, we can update the estimators as

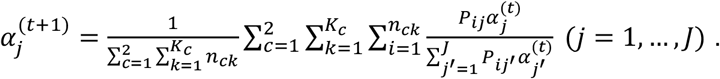

We use the above algorithm to update estimated PAS proportions until convergence.

Some genes have a large number of annotated PASs in PolyA_DB (Figure S1). The above EM algorithm would give most PASs non-zero proportions even though some isoforms have a very low abundance. To address this issue, MAAPER uses a forward selection procedure to select PASs into the model based on their statistical significance. MAAPER first ranks all PASs based on their supporting read numbers, and starts the selection process with the PAS with the largest read number. At each subsequent step, the likelihood ratio test is carried out to determine if addition of a PAS would significantly lead to increase of the likelihood. The process is terminated when no such an increase can be achieved. In essence, MAAPER selects a minimal set of annotated PASs that can sufficiently explain the observed nearSite reads. Our previous work has shown that a similar stepwise selection procedure is very efficient in identifying truly expressed RNA isoforms from RNA-seq data (Li et al., 2019).

### Unpaired statistical test for differential APA profiles

With the predicted PASs, MAAPER analyzes differential APA isoform abundances between the two conditions. Suppose there are *J* PASs in a given gene. We denote their proportions in condition *c* (*c* = 1 or 2) as 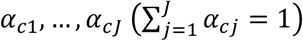. Following the notations described above, the joint likelihood of nearSite reads mapped to the gene can be derived as

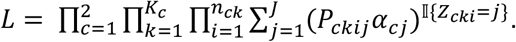

Unlike tools that focus on proximal and distal PASs only, we designed a likelihood ratio test to detect differential abundances for all PASs. The null hypothesis is *H*_0_: *α*_1*j*_ = *α*_2*j*_ (*j* = 1, …, *J*); the alternative hypothesis is *H* _*α*_: ∃ *j* ∈ {1, …, *J*} s. t. *α*_1*j*_ ≠ *α*_2*j*_. Under *H*_0_, the proportions can be estimated using the EM algorithm described above, and we denote the estimated values after convergence as 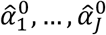. Under *H*_*a*_, we can also use an EM algorithm to update the estimators using the following rule:

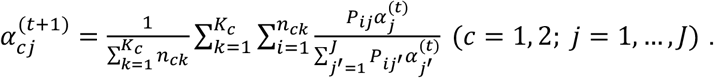

We denote the estimated values after convergence as 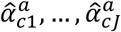. Under *H*_0_, the statistic below follows a chi-square distribution:

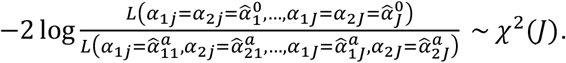

The *p* values of the likelihood ratio tests can then be obtained based on the above test statistic, and are adjusted by the Benjamini and Hochberg method (Benjamini and Hochberg, 1995) to correct for the multiple testing issue and derive false discovery rate (FDR). In our analysis, we use the adjusted *p* value = 0.01 as the threshold for identifying genes with significantly differential APA isoform abundances. In addition to the likelihood ratio test, MAAPER also provides the Fisher’s exact test to examine if the differences are significant for the two PASs with the largest proportion changes. When there are replicates, reads are first pooled from the replicates to carry out the Fisher’s test.

### Paired statistical test for differential APA profiles

In scRNA-seq experiments, the comparison of APA profiles is often among the same group of subjects. For example, it is often of interest to test for differential APA expression between two cell types from the same subjects, or between pre- and post-treatment cells from the same subjects. To address this type of questions, we propose a test for differential APA profiles among paired subjects. Suppose there are *J* predicted PASs in a given gene. We denote their proportions in sample *i* (*i* = 1,2, …, *n*) in condition *c* (*c* = 1 or 2) as 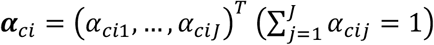 . The reads mapped to the given gene in each nearSite read sample are assigned to the PASs based on the read-PAS probabilities calculated in the prediction stage. We denote the read counts as ***X***_1*i*_ ∈ N^*J*^ and ***X***_2*i*_ ∈ N^*J*^, where *X*_*ciJ*_ is the read count of PAS *j* in sample *i* in condition *c*.

We assume that both ***X***_1*i*_ and ***X***_2*i*_ follow a Multinomial distribution

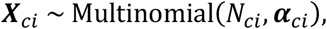

where 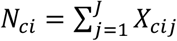. In addition, we assume that the PAS proportions in samples of the same condition independently follow the same distribution, where

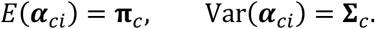

To account for the correlation between APA profiles of the same subject, we further assume that

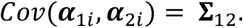

It follows from the above assumptions that

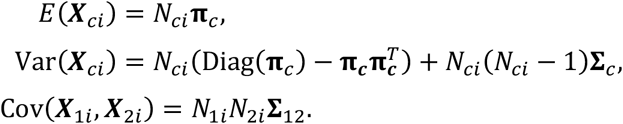

In out paired test, the null hypothesis is *H*_0_: *π*_1*j*_ = *π*_2*j*_ (*j* = 1, …, *J*); the alternative hypothesis is *H*_*a*_: ∃ *j* ∈ {1, …, *J*} s. t. *π*_1*j*_ ≠ *π*_2*j*_. Based on (Shi and Li, 2017), we can construct the following *F* statistic to test the hypotheses

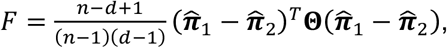

where Θ is the Moore-Penrose pseudoinverse of 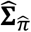 .

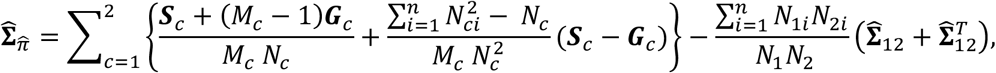

where 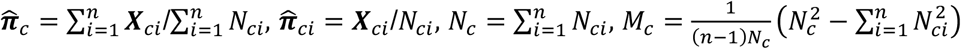, and

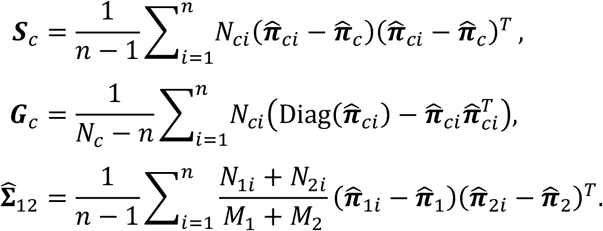

Under *H*_0_, the *F* statistic follows an asymptotic *F*-distribution with degrees with freedom *J* - 1 and *n* - *J* - 1. The *p* values can then be obtained based on the above test statistic, and are adjusted by the Benjamini and Hochberg method to correct for the multiple testing issue.

### APA regulation scores

MAAPER calculates three scores for APA regulation, namely relative length difference (RLD) for 3’UTR size difference (RLD*u*, considering 3’-most exon PASs only), RLD for pre-mRNA size difference (RLD*i*, considering intronic PASs), and relative expression difference (RED, considering all PASs).

For RLD*u*, where we only consider 3’UTR PASs, suppose a gene has *J* 3’UTR PASs, ordered from 5’ to 3’ on chromosome. Their proportions are denoted as *α*_11_, …, *α*_1*J*_ in condition 1 and *α*_21_, …, *α*_2*J*_ in condition 2. We use *l*_&_ to denote the relative 3’ UTR length of the *j*-th PAS, with *l* = 1 nt for the 5’-most PAS and *l*_*j*_ (*j* > 1) is the nucleotide distance between the first and the *j*-th PASs on the 3’UTR. The RLD*u* score between conditions A and B is calculated as 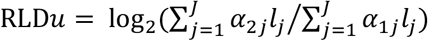 . As such, RLD*u* measures the relative length of 3’ UTR between the two conditions.

For RLD*i*, where we consider PASs located in introns and internal exons in addition to 3’-most exons, suppose a gene has *J* PASs, ordered from 5’ to 3’ on chromosome. *J*_2_ is the set of PASs that are located in introns or internal exons and *J*_(_ is the set of PASs in the 3’-most exon (*J*_1_+ *J*_2_ = *J*). *l*_*j*_ denotes the length of the pre-mRNA transcript for the *j*-th PAS, i.e, the genomic distance between transcription start site and the *j*-th PAS. Then the RLD*i* score can be calculated as 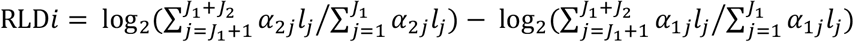. As such, RLD*i* measures the relative length of pre-mRNA transcripts with internal polyadenylation between the two conditions.

For the RED (relative expression difference) score, we use the two PASs with the largest proportion changes between two conditions. We refer to the two PASs as the distal PAS and proximal PAS based on their chromosomal positions relative to the transcription start site. Suppose the proportions of the proximal PAS in conditions A and B are denoted as *α*_$F_ and *α*_(F_, and the proportions of the distal PAS are denoted as *α*_1*p*_ and *α*_2*p*_. The RED score between conditions A and B is calculated as RED = log_2_ (*α*_2*d*_ /*α*_2*p*_) – log_2_(*α*_1*d*_/*α*_1*p*_). Therefore, a negative RED value indicates transcript shortening and a positive value transcript lengthening.

### QuantSeq data analysis

The processing of QuantSeq data followed the protocol on the website of Lexogen (https://www.lexogen.com). Briefly, the adapter contamination, polyA read through, and low quality tails were first trimmed using bbmap(Bushnell, 2014). The cleaned reads were then mapped to the RefSeq mm9 genome using STAR (Dobin et al., 2013) with the parameters of “--outFilterType BySJout --outFilterMultimapNmax 20 --alignSJoverhangMin 8 --alignSJDBoverhangMin 1 --outFilterMismatchNmax 999 --outFilterMismatchNoverLmax 0.1 --alignIntronMin 20 --alignIntronMax 1000000 --alignMatesGapMax 1000000 --outSAMattributes NH HI NM MD”. MAPPER was applied to aligned QuantSeq FWD reads from NT, AS, RC4, and RC8 samples separately for PAS prediction and differential APA analysis. Only genes with at least 50 mapped reads were considered. At the PAS prediction stage, a proportion of 0.05 was used as the threshold to select PASs. We evaluated the accuracy of MAAPER by varying this threshold between 1% and 10%, and found that MAAPER’s precision was consistently above 85% and recall above 80%, demonstrating robustness of MAAPER (Figure S16). At the APA analysis stage, an FDR-adjusted *p* value of 0.01 was used as the threshold to select significant cases. For PAS identification with QuantSeq REV data, the PAS positions were directly determined as the 3’-most positions of mapped reads.

### scRNA-seq data analysis

10x Genomics scRNA-seq data of placental cells were generated by Vento-tormo et al. (Vento-Tormo et al., 2018), which contained five donors (D8, D9, D10, D11, and D12). Reads were aligned against the hg19 human reference genome using the Cell Ranger Single-Cell Software Suite (version 3.0, 10x Genomics). Reads from three cell types (VCTs, EVTS, and SCTs) were individually aggregated into pseudo-bulk bam files for each donor. Cell type annotations were obtained from Vento-tormo et al. (Vento-Tormo et al., 2018). MAPPER was then applied to the pseudo-bulk bam files for PAS prediction and differential APA analysis. Same thresholds were used as in the QuantSeq FWD data analysis. Data representation by tSNE based on gene expression levels was carried out by the R package Seurat (version 3.1.2) (Satija et al., 2015).

## Data Availability

QuantSeq FWD and REV datasets generated in this study have been deposited into the GEO database under the accession number GSE164958.

## Software Availability

The MAAPER method has been implemented as an R package, which is available at Github (https://github.com/Vivianstats/MAAPER).

## Author’s contributions

WVL and BT conceived the analyses. DZ and BT designed the QuantSeq experiments and DZ performed the experiments. WVL and BT developed the method and WVL implemented the method and software. WVL and RW analyzed the data. WVL and BT wrote the paper.

## Acknowledgements

We thank Chuwei Zhong and other members of BT lab for helpful discussions. Computational resources were provided by the Office of Advanced Research Computing (OARC) at Rutgers, The State University of New Jersey.

