## Supplementary File for "MAAPER: model-based analysis of alternative polyadenylation using 3’ end-linked reads"

**Table S1. Read number of QuantSeq samples.**

| Sample | Number of QuantSeq FWD reads | Number of QuantSeq REV reads |
|---|---:|---:|
| NT | 27,859,996 | 1,817,267 |
| AS | 27,859,632 | 3,928,573 |
| RC4 | 28,788,156 | 1,529,184 |
| RC8 | 27,948,416 | 3,605,416 |

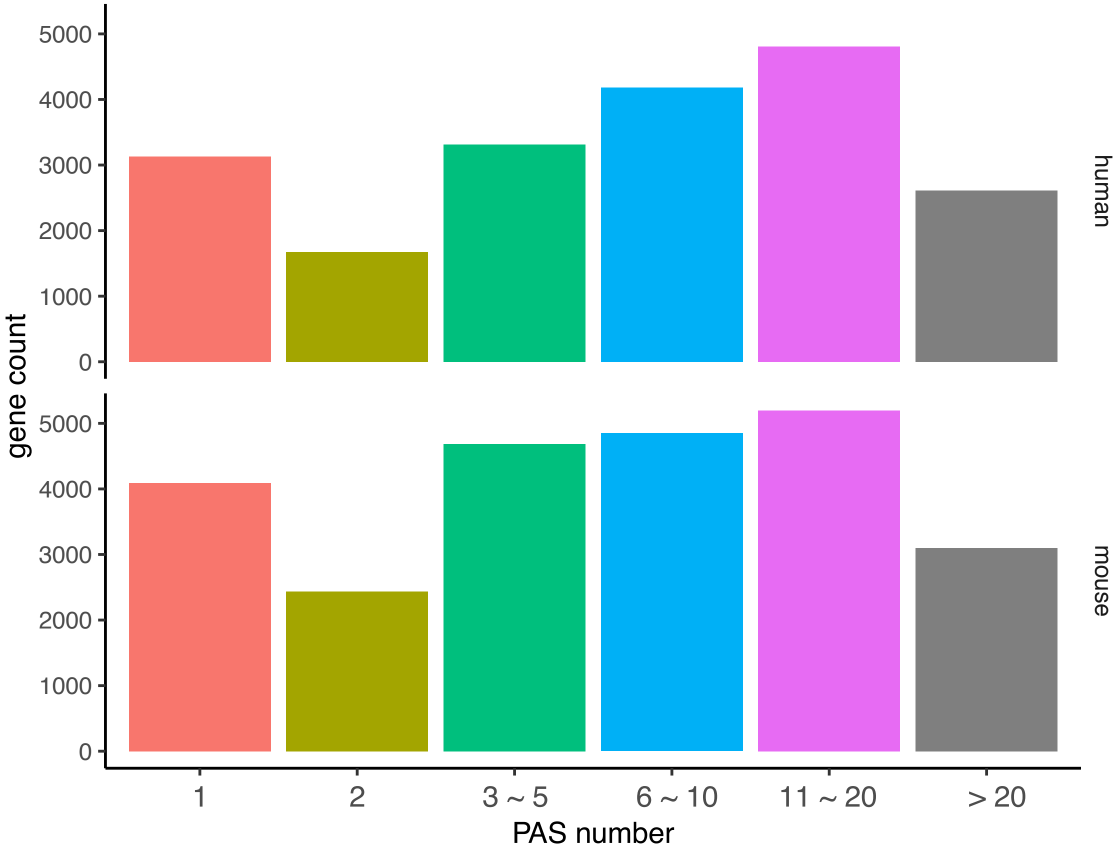


**Figure S1**. Number of annotated PASs in PolyA_DB.
The numbers of human and mouse genes with varying number of annotated PASs are displayed as barplots. The PAS number of human genes has a median of 7 and a mean of 10.1. The PAS number of mouse genes has a median of 6 and a mean of 9.7.

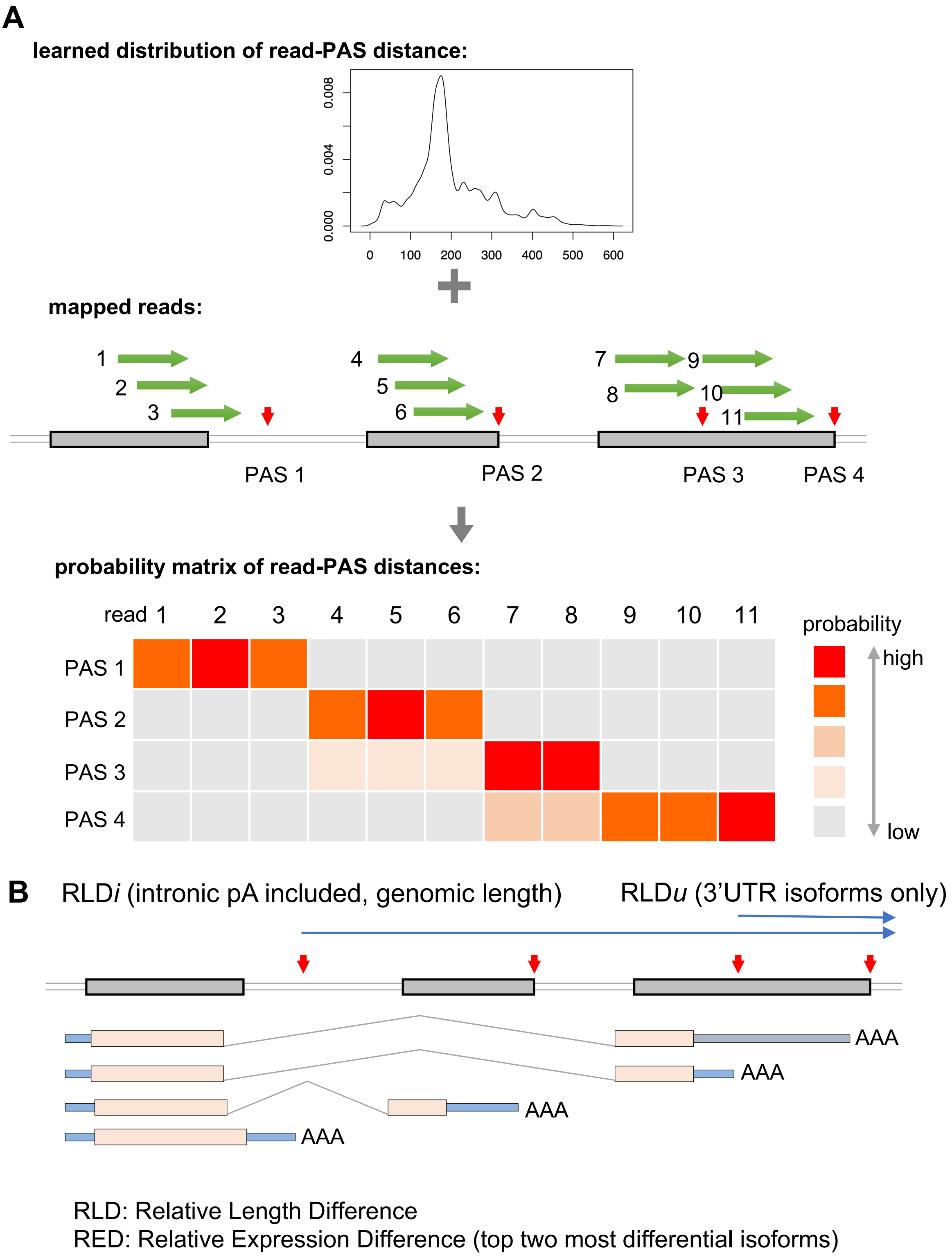


**Figure S2**. Schematic illustrations of MAAPER.
**A**: Illustration of the calculation of probabilities of read-PAS distances. **B**: Illustration of the RLD and RED scores.

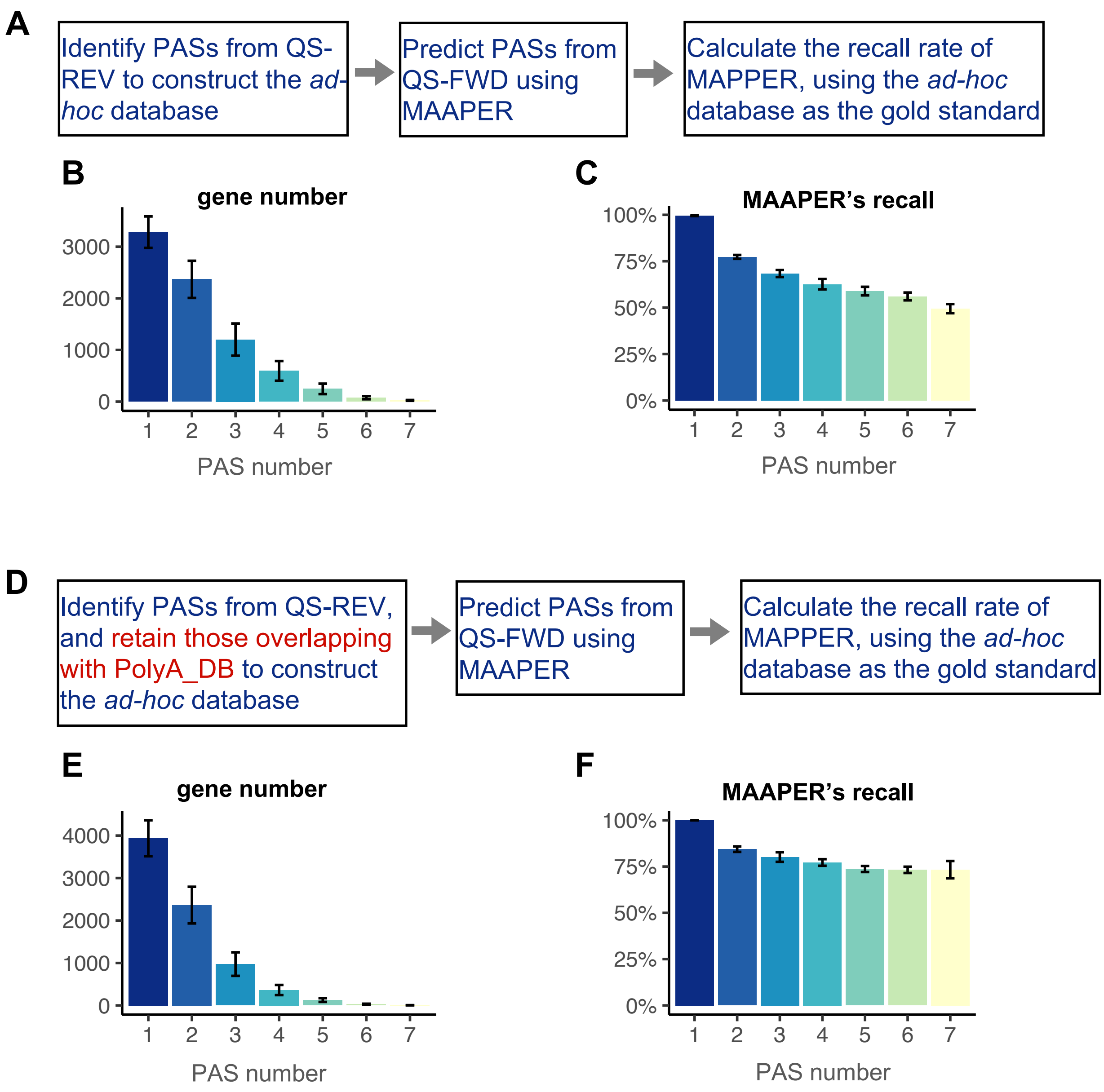


**Figure S3**. Recall rates of MAAPER in PAS prediction.
**A-C**: Results are based on identified PASs from QS-REV data. **D-F**: Results are based on identified PASs (from QS-REV data) that are also annotated in PolyA_DB. **A,D**: Schematic diagrams showing steps to calculate MAAPER's recall rate. **B,E**: Number of genes with different number of identified PASs from QS-REV data. Gene number is averaged across NT and AS-treated conditions, and error bars represent one standard deviation. **C,F**: Recall rates of MAAPER's prediction from QS-FWD data for genes with varying number of identified PASs from QS-REV data. Recall rate is averaged across NT and AS-treated conditions, and error bars represent one standard deviation.

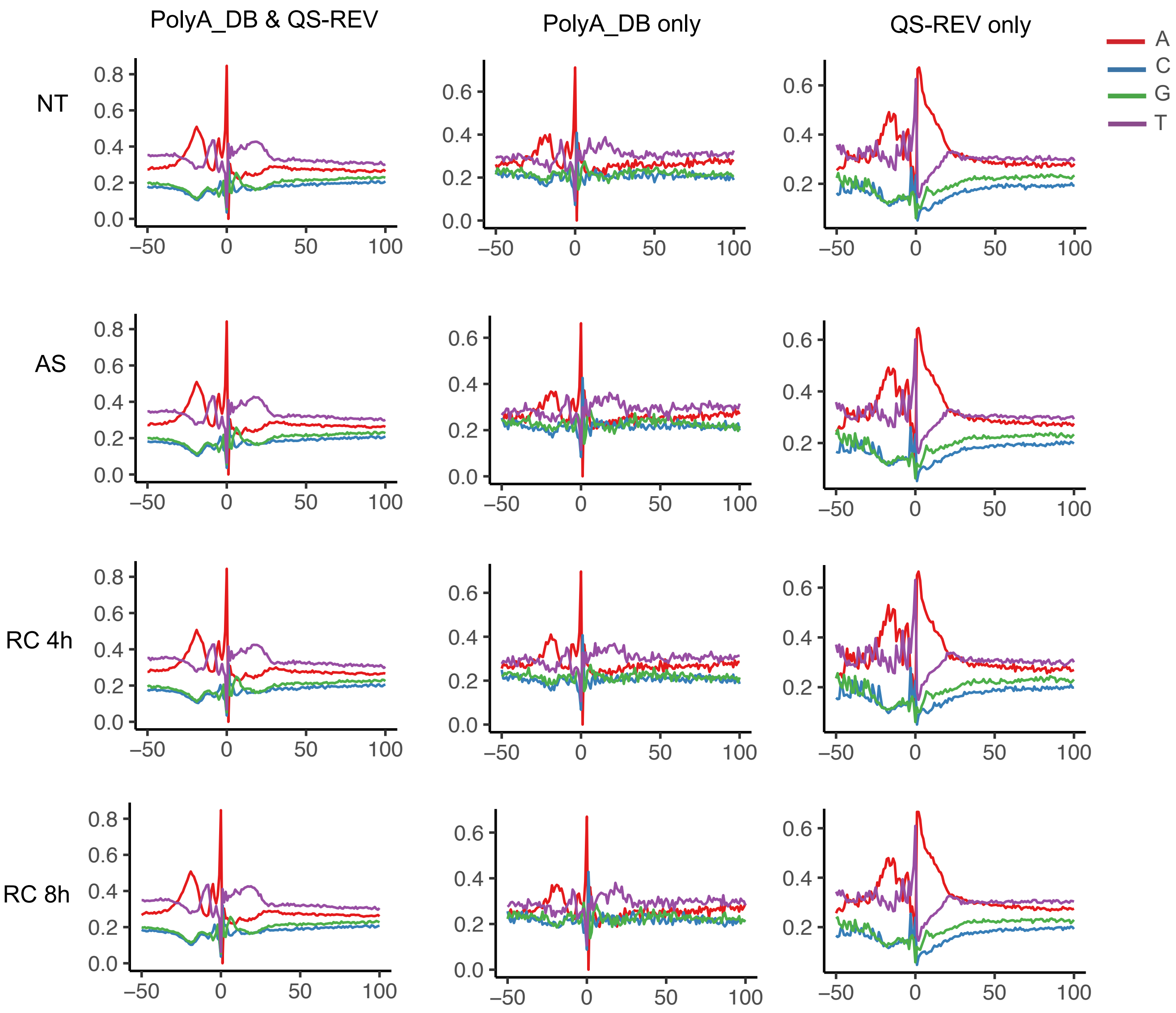


**Figure S4**. Nucleotide frequency around PASs.
QS-FWD reads were assigned to three types of PASs: PASs predicted by MAAPER, annotated in PolyA_DB, and supported by QS-REV reads; PASs predicted by MAAPER, annotated in PolyA_DB, but not supported by QS-REV reads; PASs not annotated in PolyA_DB but supported by QS-REV reads. The average nucleotide frequency in these three types of PASs were calculated for each sample.

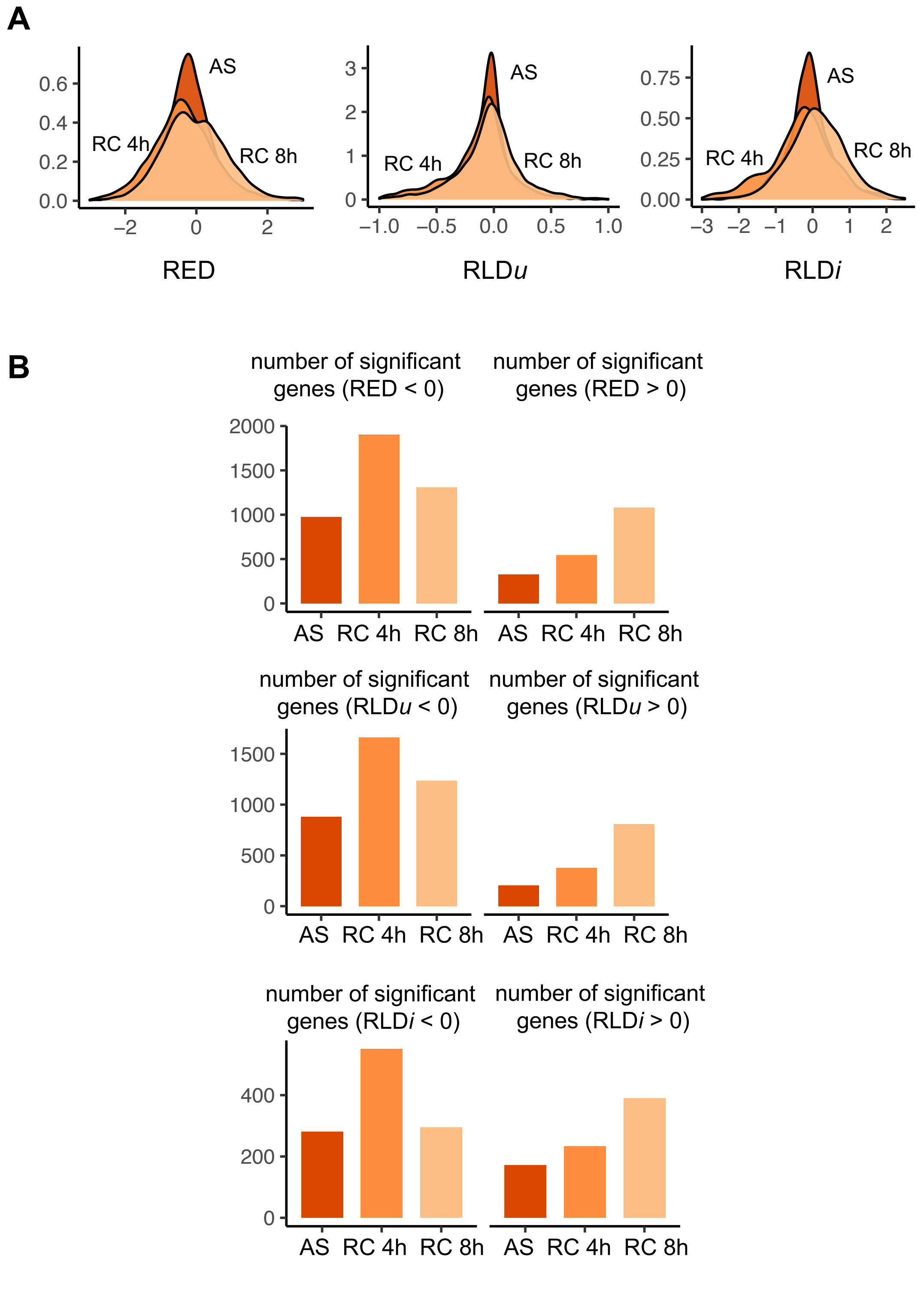


**Figure S5**. RED and RLD scores for AS data.
**A**: Distribution of RED, RLDu, and RLDi scores in the AS, RC 4h, and RC 8h samples. **B**: Number of genes with significant APA changes (FDR-adjusted *p* value < 0.01) with negative or positive RED/RLD scores in each sample.

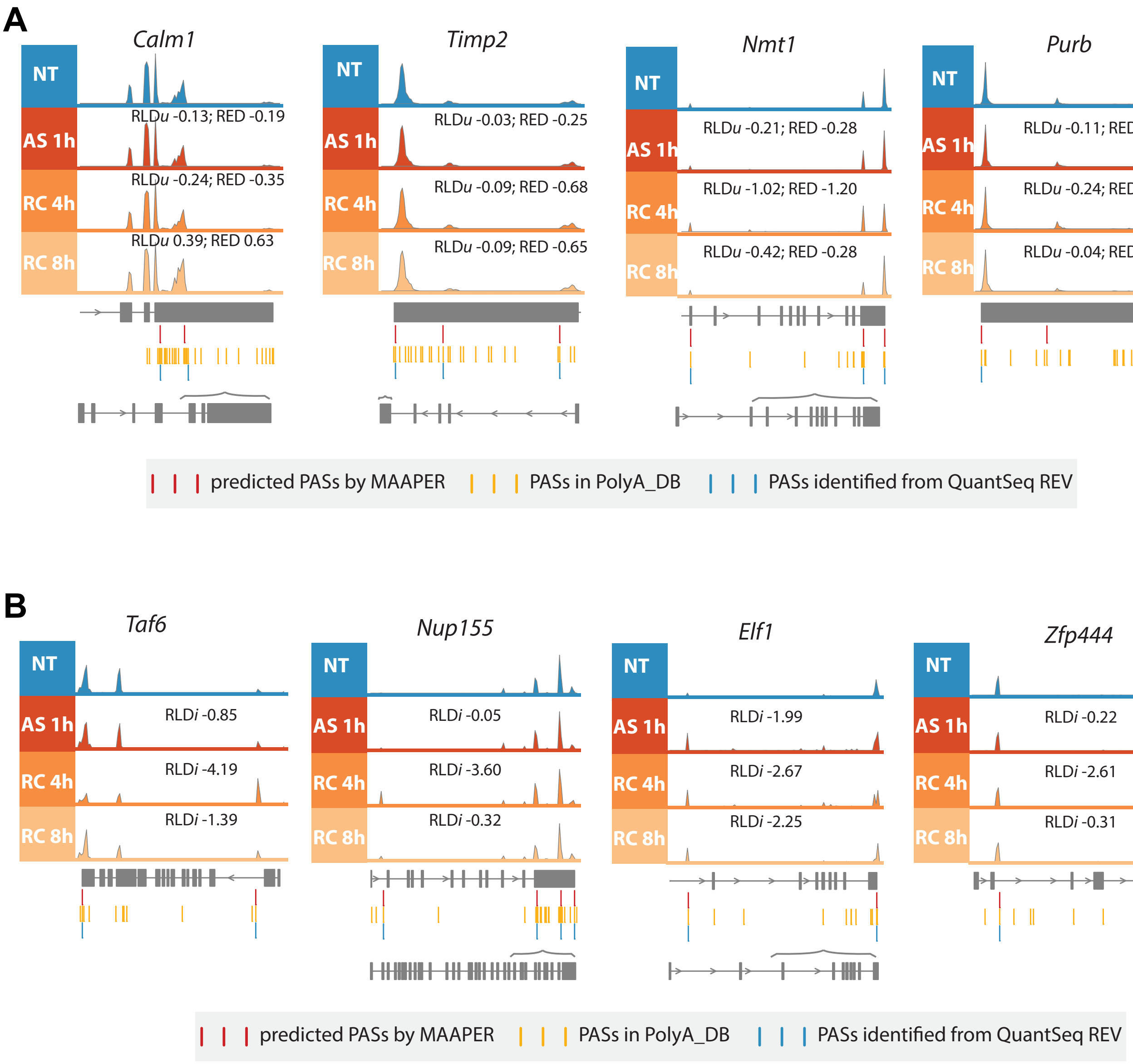


**Figure S6**. Read alignment tracks of example genes in NT and AS-treated samples. **A**: Four genes whose 3'UTR shortening has been validated by PCR experiments. **B**: Four genes with significant activation of intronic PASs in AS-treated conditions.

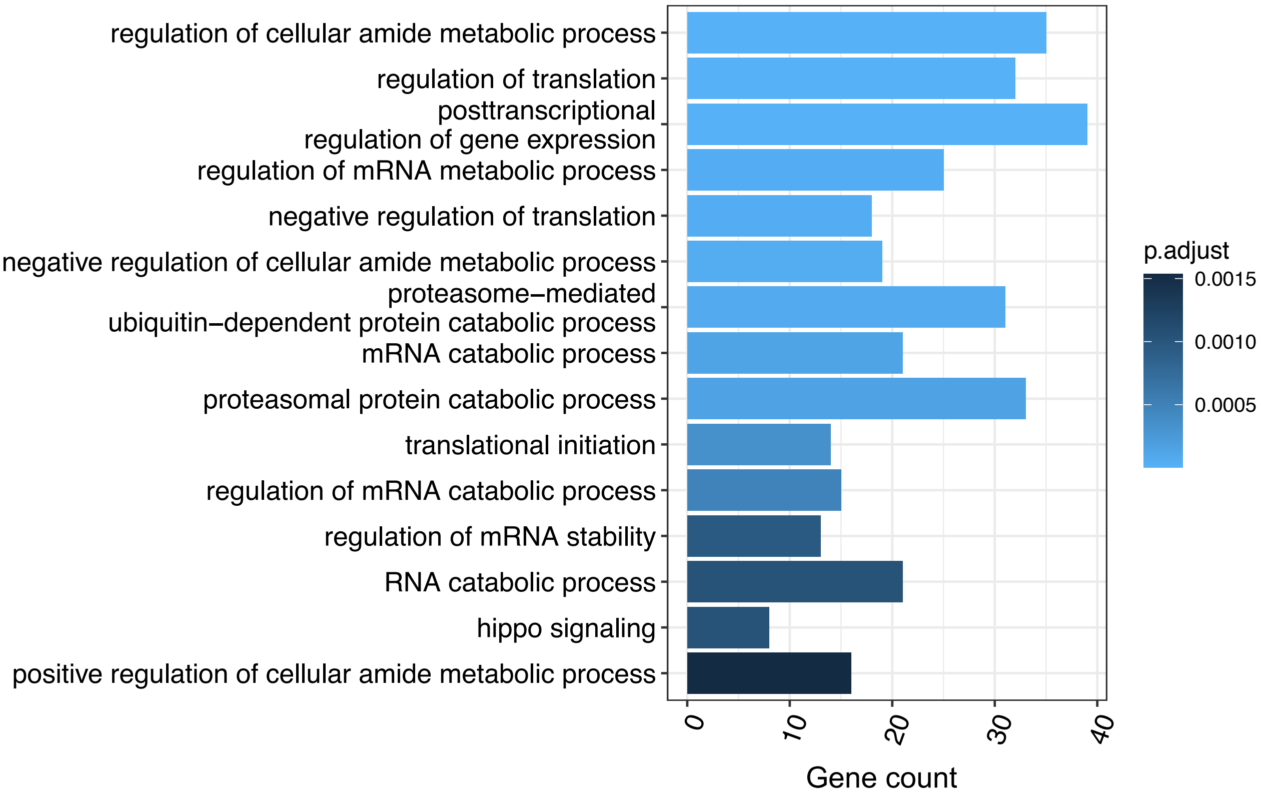


**Figure S7**. Top 15 enriched GO terms of biological processes in genes with 3'UTR shortening. Genes with significant APA changes and negative RLD*u* scores were selected and the 15 GO terms with the smallest FDR-adjusted *p* values are displayed.

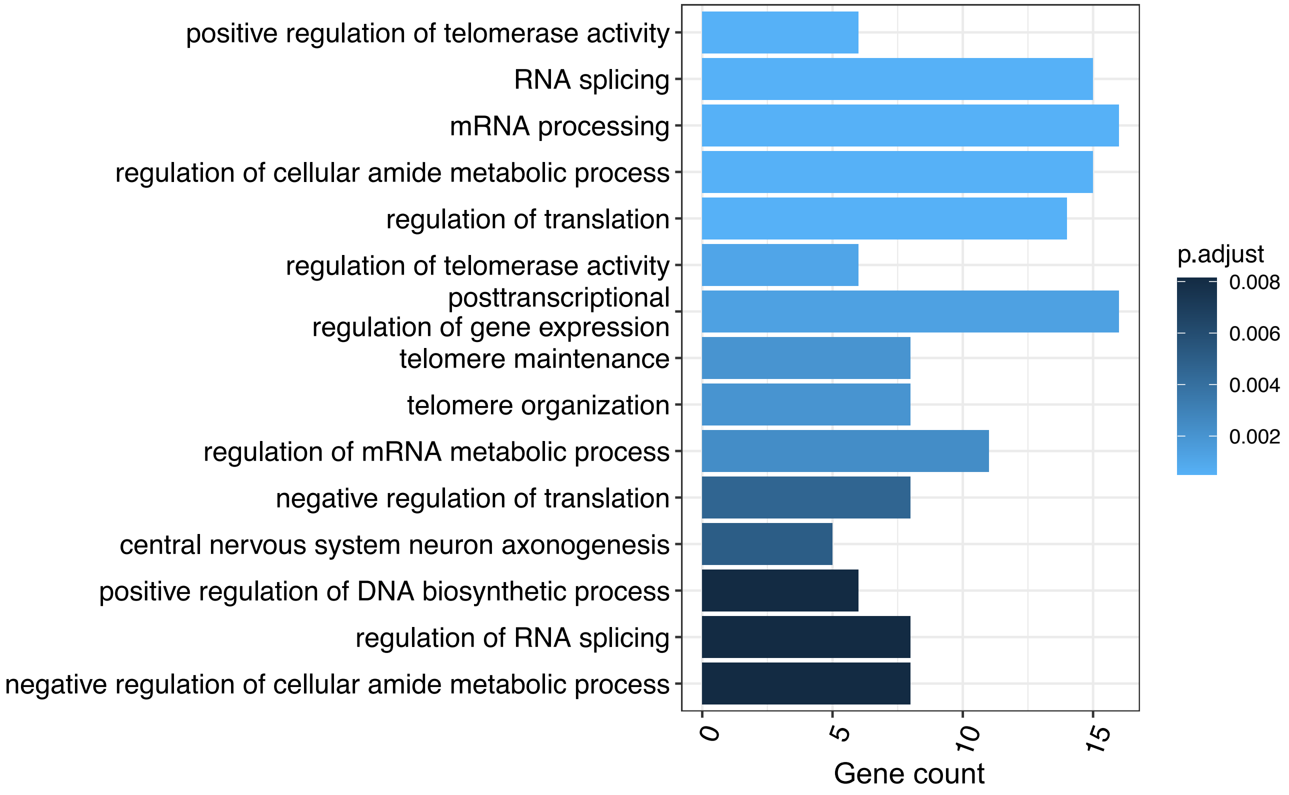


**Figure S8**. Top 15 enriched GO terms of biological processes in genes with pre-mRNA shortening. Genes with significant APA changes and negative RLD*i* scores were selected and the 15 GO terms with the smallest FDR-adjusted *p* values are displayed.

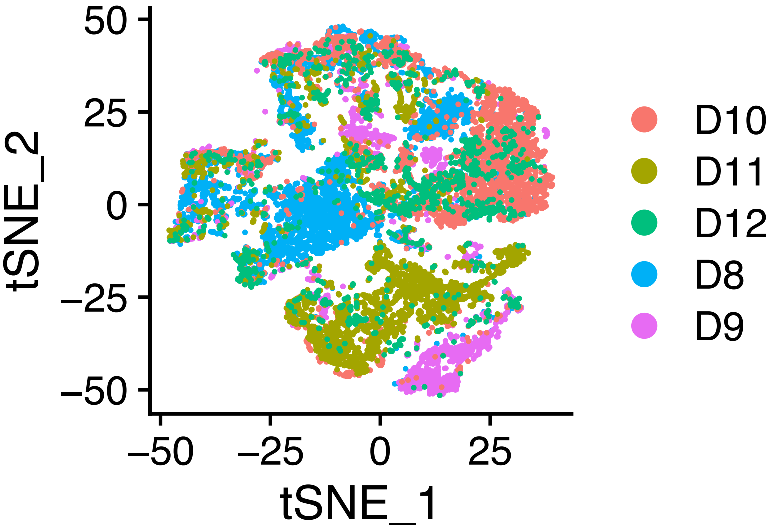


**Figure S9**. tSNE plots of the trophablast cells from the five donors.

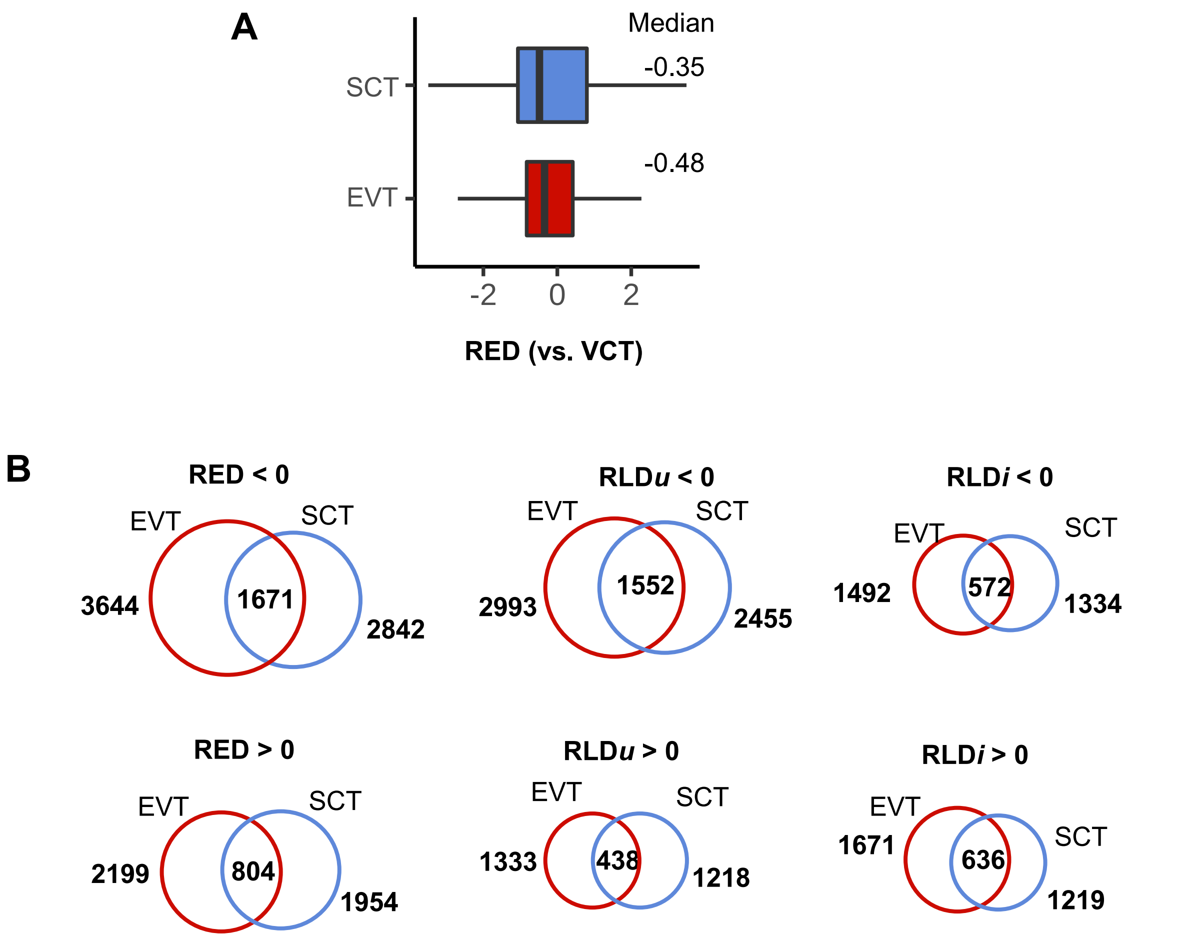


**Figure S10**. RED and RLD scores for comparison between SCTs or EVTs and VCTs. **A**: Boxplots of RED scores calculated for EVTs and SCTs, based on genes that showed significant APA changes in each cell type. Median values are labelled on the bottom. **B**: Venn diagrams showing the overlap of genes with significant APA changes (FDR-adjusted *p* value < 0.01) in SCTs and EVTs.

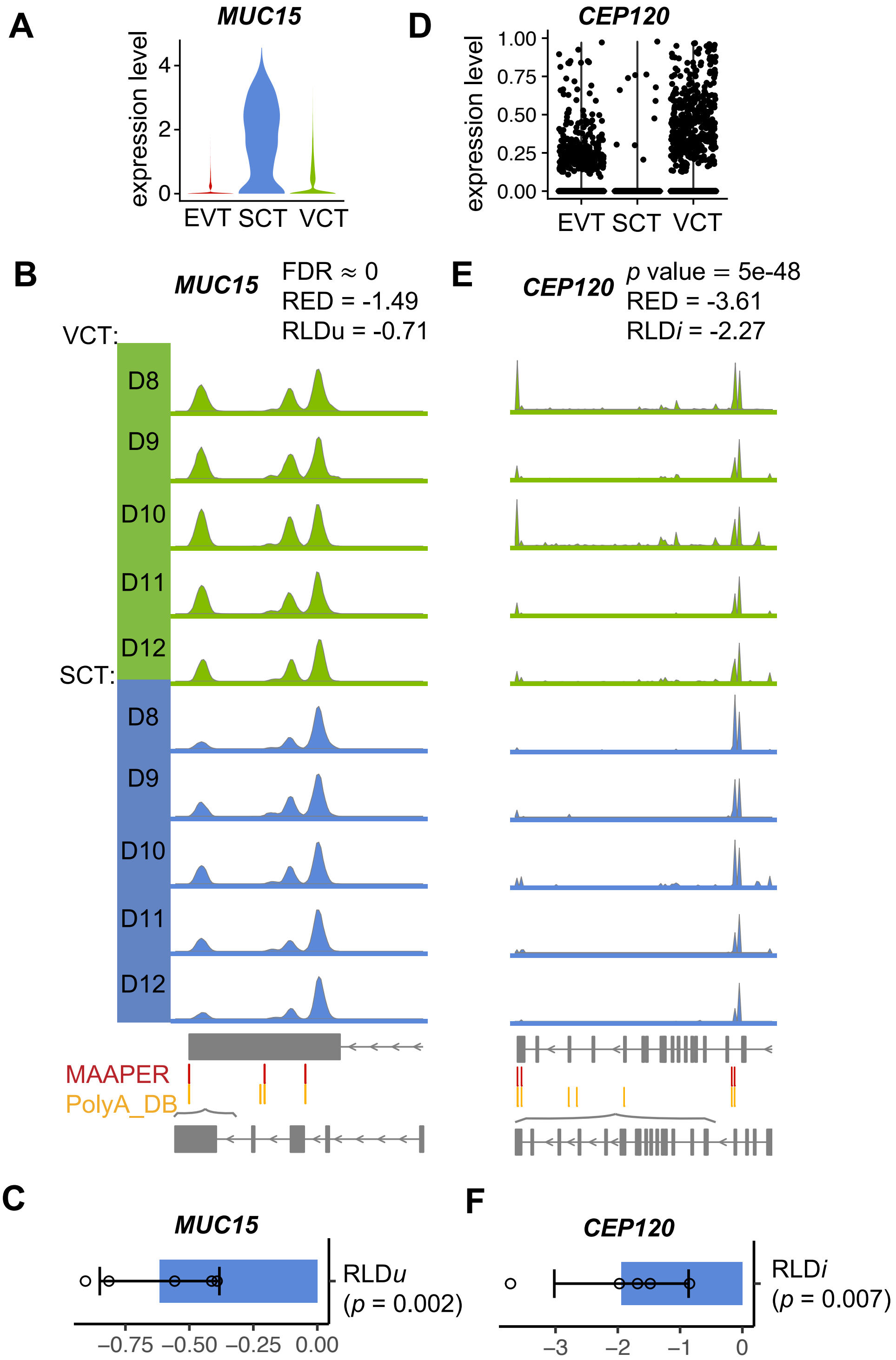


**Figure S11**. (**A**,**D**): Log10 transformed gene expression of *MUC15* and *CEP120* in the three cell types. (**B**,**E**): Read alignment tracks of the *MUC15* and *CEP120* genes in VCT and SCT cells. (**C**): RLD*u* scores of *MUC15* in the five donors. (**F**): RLD*i* scores of *CEP120* in the five donors.

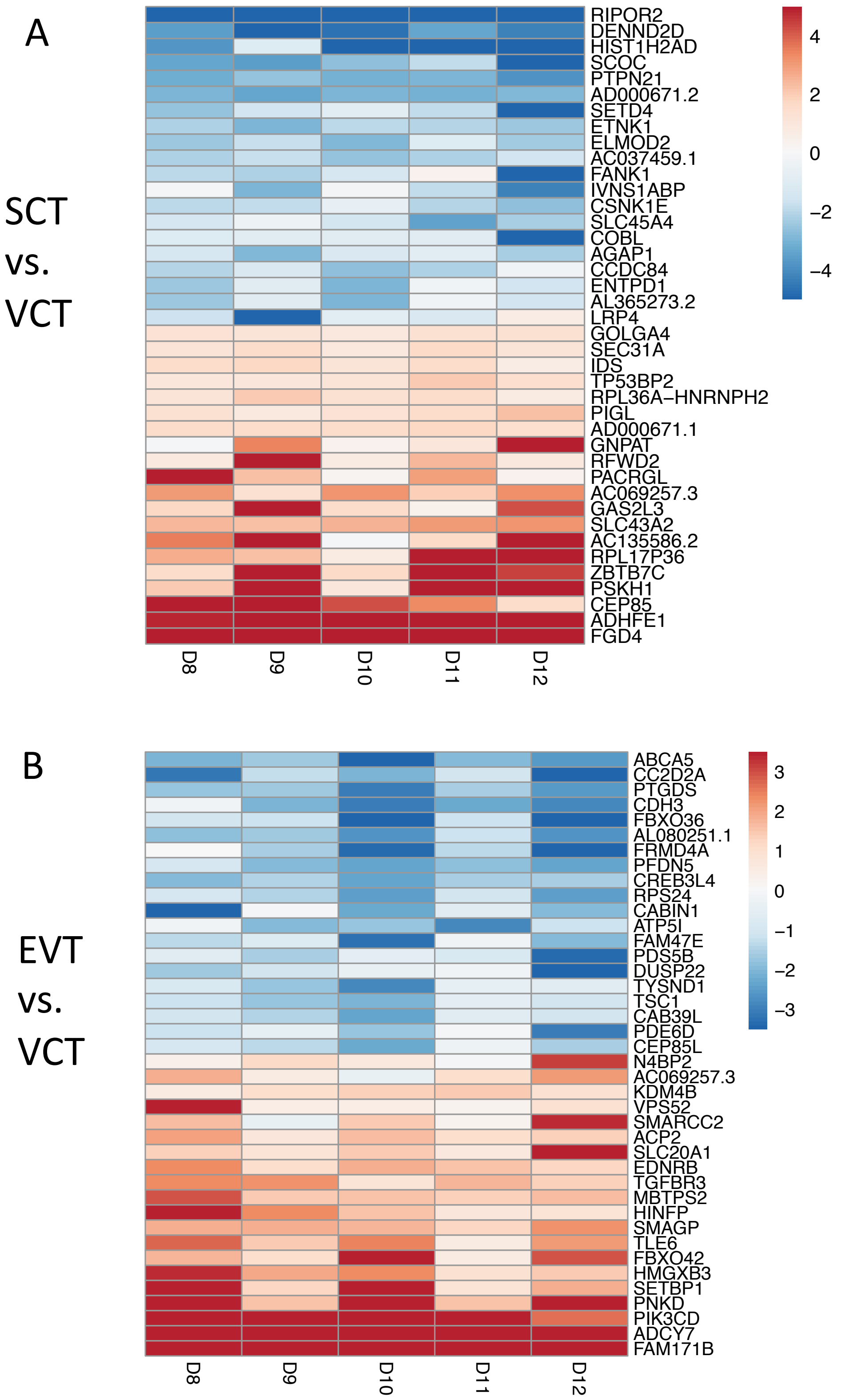


**Figure S12**. Genes with significant APA changes identified by the paired test. (**A**): RED scores for comparison between SCTs and VCTs. (**B**): RED scores for comparison between EVTs and VCTs. Top 40 genes with significant p values and largest absolute values of mean RED are displayed.

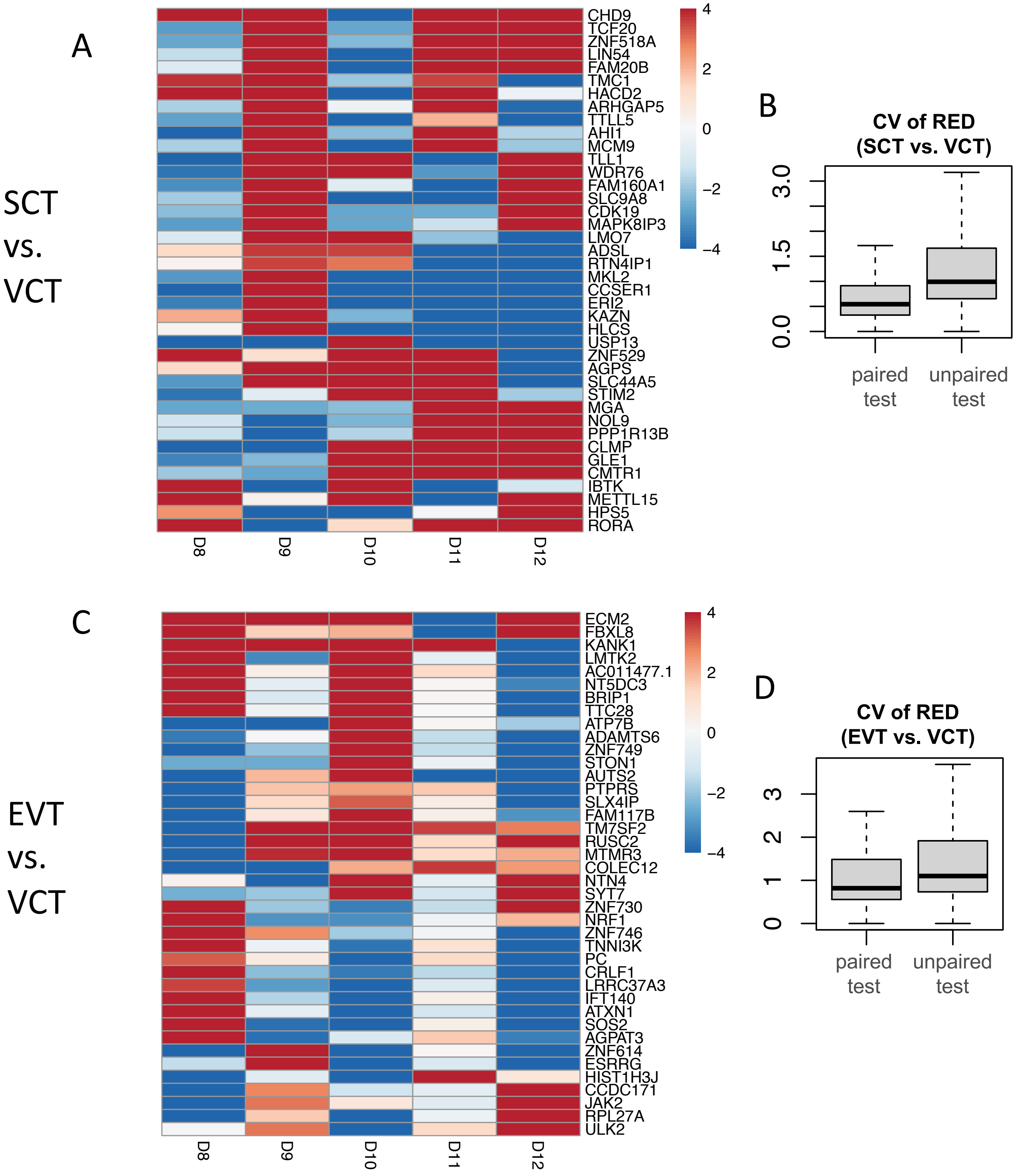


**Figure S14**. Genes with significant APA changes identified by the unpaired test. (**A,C**): RED scores for comparison between SCTs (A) or EVTs (C) and VCTs. Top 40 genes with significant p values and largest variation of RED are displayed. (**B,D**): Coefficient of variation (CV) of significant genes' RED scores.

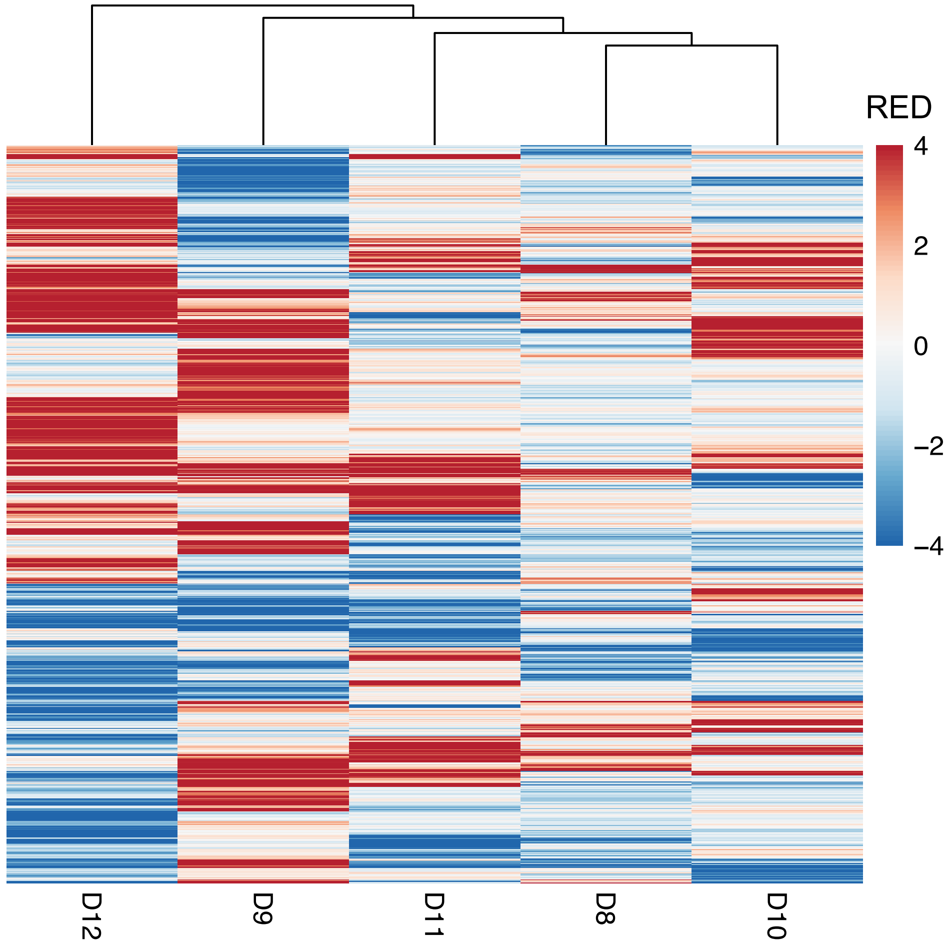


**Figure S15**. Hierarchical clustering of the five donors based on genes with highly variable RED scores (SD > 1.5).

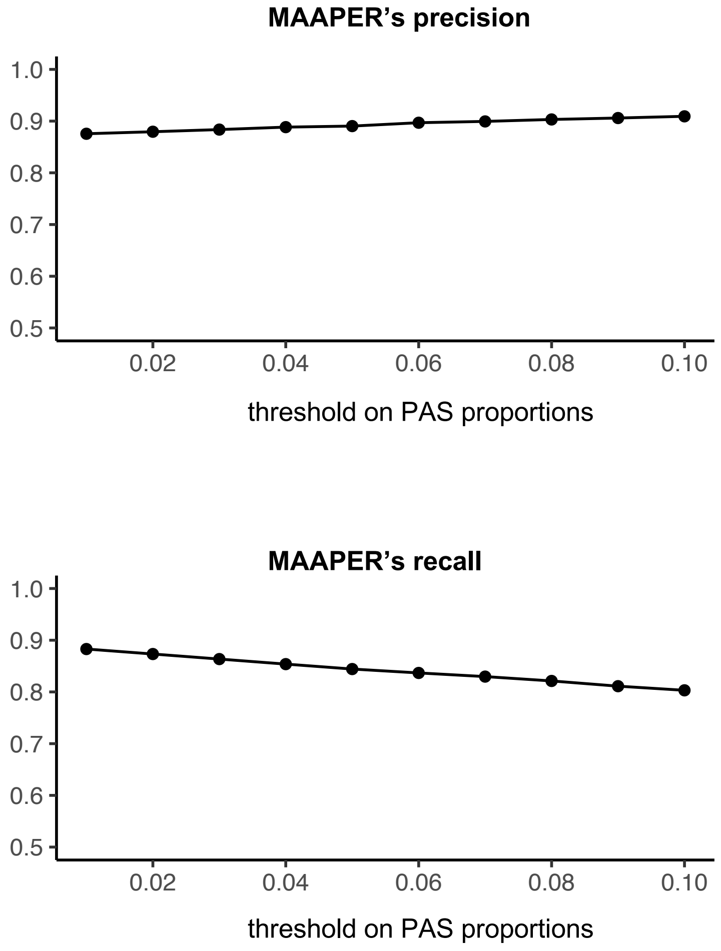


**Figure S16**. Sensitivity analysis of MAAPER.
Precision and recall rates were calculated by comparing MAAPER's results with the identified PASs (restricted to those overlapping with PolyA_DB) from QuantSeq REV data. MAAPER's threshold on PAS proportions varied between 1% and 10%. Analysis was performed using the RC4 sample as an example.
